# On demand nanoliter sampling probe for collection of brain fluid

**DOI:** 10.1101/2022.04.08.487549

**Authors:** Joan Teixidor, Salvatore Novello, Daniel Ortiz, Laure Menin, Hilal A. Lashuel, Arnaud Bertsch, Philippe Renaud

## Abstract

Continuous fluidic sampling systems allow collection of brain biomarkers *in vivo*. Here, we propose a new sampling paradigm, Droplet on Demand (DoD), implemented in a microfabricated neural probe. It allows sampling droplets loaded with molecules from the brain extracellular fluid punctually, without the long transient equilibration periods typical of continuous methods. It uses an accurate fluidic sequence and correct operation is verified by the embedded electrodes. As a proof of concept, we demonstrated the application of this novel approach *in vitro* and *in vivo*, to collect glucose in the brain of mice, with a temporal resolution of 1-2 minutes and without transient regime. Absolute quantification of the glucose level in the samples was performed by direct infusion nanoelectrospray ionization Fourier transform mass spectrometry (nanoESI-FTMS). By adjusting the diffusion time and the perfusion volume of DoD, the fraction of molecules recovered in the samples can be tuned to mirror the tissue concentration at accurate points in time. This makes quantification of biomarkers in the brain possible within acute experiments of only 20 to 120 minutes. DoD provides a complementary tool to continuous microdialysis and push-pull sampling probes. The advances allowed by DoD will benefit quantitative molecular studies in the brain, namely for molecules involved in volume transmission or for protein aggregates that form in neurodegenerative diseases over long periods.

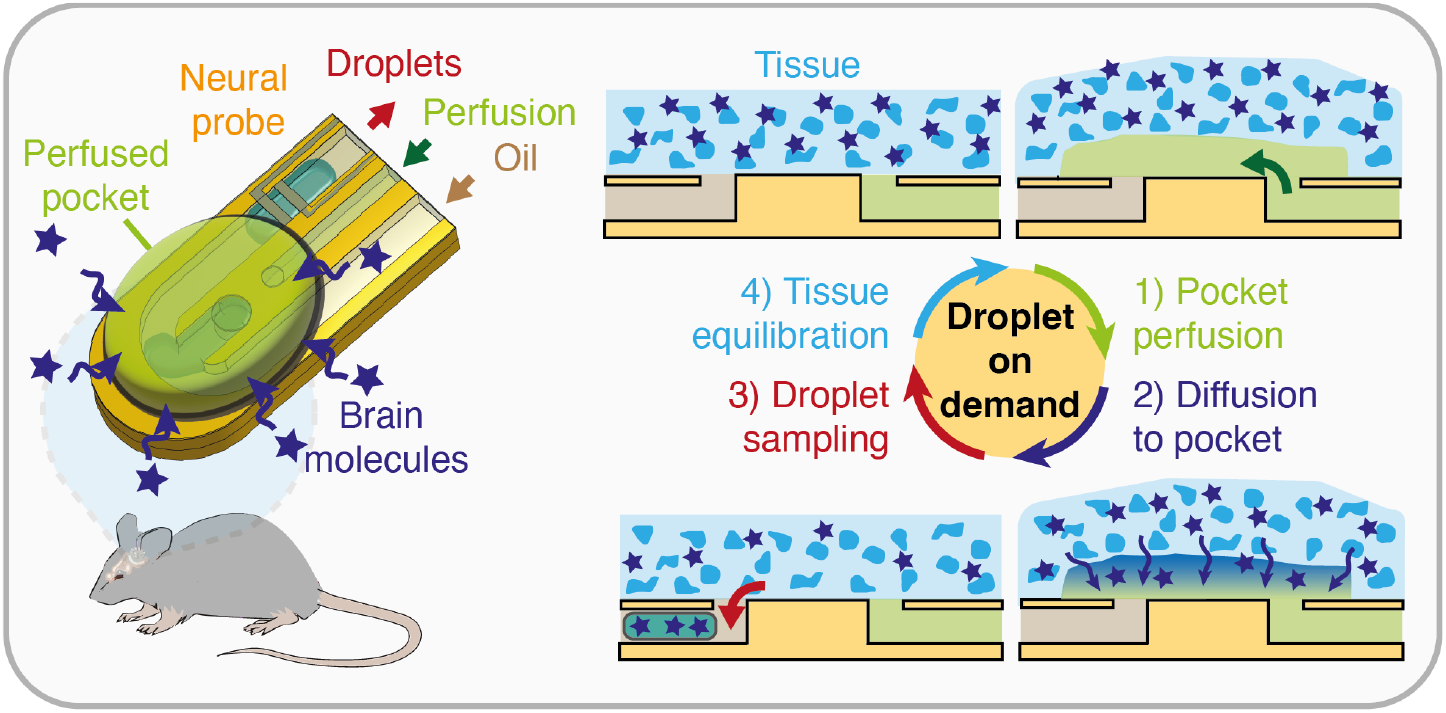

## 1. INTRODUCTION

Minimally invasive solutions to study molecular processes in the brain *in vivo* mostly rely on electrochemical and optical spectroscopy systems, as well as on fluidic sampling probes coupled to analytical systems^1^. The first two types of systems present high spatial and temporal resolutions, and probe the molecules directly in the tissue, based on charge transfer processes and light-matter interactions, respectively. Small electrodes are particularly suited for electrical stimulation^2^ and sensing, to resolve fast signals at the sub-cellular level^3^, to map the interconnections between neurons^4^ or to correlate electrical and chemical events^5^. Alternatively, fluidic sampling systems such as microdialysis and push-pull probes, enable studying the interplay of multiple molecules in parallel, captured by diffusion from the brain extracellular fluid (ECF) into a perfusion fluid. Being compatible with a wide range of analytical methods, off-line analysis of a large panel of molecules is possible. However, reducing their footprint is limited by the hydraulic resistance and the risk of clogging of small channels. Most invasive systems have benefited from technological developments in material sciences and microfabrication technologies. They allowed integrating new stimulation and sensing functions (implantable neurostimulation probes^6^, electronic dura mater with drug delivery^7^, fiber-based multimodal probes^8^) and new materials (glassy carbon electrodes in polyimide probes for dopamine measurement^9^, optofluidic probes with µLED for optogenetics^10^), while reducing their footprint. This benefited their performances and extended the possible *in vivo* studies. Review articles compared these technologies^1,11–14^ and some non-invasive imaging methods^15,16^.

Among the fluidic sampling probes, the microdialysis probe is a gold standard. It possesses two capillaries surrounded by a semi-permeable membrane at the tip, that confines the flow of perfusion fluid, called perfusate, within the probe on one side and contacts the tissue on the other side^12^. The tip is continuously infused with perfusate, that collects analytes by diffusion from the tissue, through the membrane and to the perfusate, before exiting at the outlet as the dialysate. In microdialysis probes, the long membrane defines the cut-off size of the collected molecules and the perfusate/tissue interface, but limits the spatial resolution. This is improved in push-pull probes, membrane-less fluidic sampling probes, that consist of two capillaries. One infuses perfusate at the tip to wet the tissue and one samples it back. The absence of membrane also enables collection of larger biomarkers. Miniaturization techniques have reduced the large footprint shared by both systems, by integrating membranes directly in polyimide^17^ and silicon probes^18^ and by scaling down the channels^19,20^. They also enabled the Kennedy group to improve the temporal resolution of these systems, which was previously limited by Taylor dispersion in the tubing, by coupling the dialysate to a droplet generator at the outlet^21^. Following these advances, new fluidic sampling probes were developed, integrating flow actuation systems^20,22^, droplet generation features^23,24^, and stimulation and sensing electrodes^5^. Our group contributed the advancements through the development of a flexible probe allowing electrostimulation, neural signal recording, and droplet sampling directly at the tip^25,26^. Following the splitting of the large volumes of dialysate into picoliter to nanoliter volume droplets, analytical methods were adapted to analyze their content and a library of highly-sensitive solutions, unperturbed by surfactants and oil^27^, was developed. This benefitted from the rich literature of droplet microfluidics (e.g., droplet generation^28^, picoinjection^29^, phase extraction^30–32^, etc.). These solutions, which were extensively covered in recent reviews^33–36^, comprise in-flow electrochemical chips^37,38^, fluorescent assays^23,39,40^ and include many developments of electrophoretic^30,41^ and mass spectrometry methods^26,42–48^.

Overall, microtechnology provided smaller probes for faster and more localized sampling, while droplets improved the temporal resolution. However, quantifying the true concentration of molecules in the tissue with continuous fluidic sampling probes remains challenging. Here, we introduce a novel sampling paradigm, Droplet on Demand (DoD), that we implemented in a microfabricated probe for acute sampling experiments, and verified through *in vitro* and *in vivo* experiments. DoD enables punctual sampling of analytes in the ECF directly in droplets and accommodates its flexible sampling sequence to the transport of molecules in the tissue. This allows collection of highly concentrated droplets mirroring the tissue concentration punctually, over a single accurate time window, or repeatedly, for quantitative molecular studies within acute experiments.

## 2. MATERIALS AND METHODS

### 2.1. Probe fabrication

The probes were built at the EPFL Center of MicroNanoTechnology, on a silicon wafer covered by a sacrificial thin film of aluminum. Supporting Information contains the details of the fabrication process (Figure S1). Briefly, a 3.5 µm layer of polyimide (PI2611, Dupont) was patterned by photolithography and plasma etching, followed by sputtering of 75-350-75 nm of titanium, platinum, and titanium. Metallic electrodes were patterned by photolithography and ion bean etching, and later encapsulated under a secondary 1 µm polyimide layer (PI2610, Dupont), patterned similarly. Fluidic channels were patterned by photolithography of a 40 µm spun coated SU-8 layer (SU-8 3025, MicroChem), followed by photolithography of a laminated 40 µm SU-8 dry film. The channels were cut open with a laser and the probes detached from the wafer by anodic release^17^. The probes were cleaned with 1% HF, deionized water, isopropanol, and ethanol and dried out with air. Fluidic channels were interfaced to 250/350 µm ID/OD fused silica capillaries (TSP-250350, BGB) with UV-sensitive epoxy. After 60 s oxygen plasma activation at 100 W and 0.4 mbar (FEMTO, Diener Electronics), microfluidic channels were incubated with 2% PFOTS (trichloro(1h,1h,2h,2h-perfluorooctyl)silane, 448931-10G, CAS 78560-45-9, Merck) in PFD (perfluoro(methyldecalin), 372439, CAS 51294-16-7, Merck) for 20 minutes at 75°C, rinsed with PFD, dried out, and incubated for 20 minutes at 75°C. Using the same activation, capillaries were incubated with Sigmacote (SL2-25mL, Merck) for 20 minutes at 20°C, dried out, and incubated for 20 minutes at 100°C. Probes were disinfected by gas phase H2O2 prior to use *in vivo*.

### 2.2. FEA Simulations

COMSOL Multiphysics 5.6 was used to model the diffusion of molecules within DoD sequences. The model considered a static parallelepipedal pocket of perfusate covering the probe, at the center of a 126 mm^3^ cube of 0.6% agarose loaded with a 1 µM glucose concentration in the entire domain. The free diffusion coefficient of glucose in the perfusate, D0, was taken as 600 µm^2^/s at 20 °C, corrected with a tortuosity^49^ factor λ of 1.6 in the agarose, defined as a porous medium. Initially, the perfusate contained 0 µM of glucose. “No flow” conditions were set on the walls of the probe and on the sides of the agarose cube. The module “transport of diluted species in porous media” was used to simulate the diffusion of glucose in the medium and in the pocket. A progressive free quadrilateral mesh was set in the pocket of perfusate, with 2 µm thick features, perpendicular to the probe surface. A progressive free tetrahedral mesh was used with features of maximum 50 µm in the 1 mm vicinity of the pocket and with coarse features further away in the rest of the medium. Simulation steps of 0.5 s were performed, with a relative tolerance of 0.005 and diffusion was the only mass transport mechanism. The CAD of the model is shown in Figure S2a. The simulations of cycles proceed as follows. The first cycle starts with a diffusion step, with analytes diffusing over tdiffusion from the tissue to the perfusate. When the pocket is sampled, it is excluded from the model and its average concentration is evaluated. Then, the equilibration step starts, and the mass transport is simulated over tequilibration, considering diffusion within the medium only. The next cycle proceeds as the first cycle, with a pocket with 0 µM of glucose, except that the new initial spatial concentration in the medium is given by the spatial concentration at the end of the equilibration step of the previous cycle. The temporal evolution of the concentration in the perfusate over 3 simulated cycles is illustrated in Figure S2b, with the domains considered at each step.

### 2.3. DoD implementation

The inert fluorinated phase PFD was degassed for 20 minutes prior to use. 300 mM sucrose (S0389, CAS 57-50-1, Merck) in deionized water and Perfusion Fluid CNS (P000151, CMA Microdialysis) were used as perfusates for *in vitro* and *in vivo* experiments respectively. Agarose 0.6% brain phantoms^50^ were prepared by dissolving agarose (A9539, CAS 9012-36-6, Merck) in tris-borate-EDTA buffer 1X (T4415, Merck). Cubes of 1 cm^3^ were cut and incubated overnight in an aqueous solution of 150 mM NaCl (S9888, CAS 7647-14-5, Merck) and 10 µM fluorescein (46955, CAS 2321-07-5, Merck). The conductivity of 0% NaCl and 100% NaCl was respectively 5.99 µS/cm and 11.02 mS/cm and increased linearly with the NaCl concentration. NeMESYS low-pressure syringe pumps (Cetoni) were used with 25 µL glass syringes (1702N, Hamilton) to drive the perfusion and inlet lines. A pressure controller Elveflow AF1 Dual (−700mbar to 1000mbar, Elvesys) connected to a flow rate sensor (MFS 2, Elvesys) drove the outlet line. Tubing consisted of rigid capillaries (TSP-250350, BGB). A LabVIEW software (Lab-VIEW 2017, National Instruments) was developed to coordinate the operation of all fluidic equipment, for timed DoD sequences, and report generation. Electrical droplet measurement and detection were performed with a HF2LI Lock-in amplifier and a HF2TA transimpedance current amplifier (Zurich Instruments), with 100 mV excitation at 1 MHz across two 20 µm wide Pt electrodes in the outlet channel. The DoD parameters used for all samplings are reported in Table S1.

### 2.4. In vivo experiments

All experimental procedures were performed in accordance with the local animal care authorities, under the cantonal license VD3492. Acute sampling experiments were performed with C57BL6/J mice. Narcosis was induced by intra-peritoneal injection (10 µL/g) of a cocktail of ketamine (100 mg/kg) and xylazine (10 mg/kg). The head of the animal was shaved, the skin cleaned with betadine and the animal placed on a stereotaxic frame to which the probe holder was secured. A lubricant eye ointment was applied to prevent dry eyes during anesthesia and a heating pad was used to control the temperature of the body during the procedure. Stereotactic coordinates were set to +0.6 mm anteroposterior, 2.0 mm mediolateral and −2.60 mm dorsoventrally from bregma, to target the dorsal striatum. A hole just big enough for the probe to access the brain was drilled into the skull without injuring the brain. To ease the insertion and target the right brain location, a 125 µm diameter optical fiber (FG105LCA, Thorlabs) was fixed on the needle of the probe. A digital endoscope enabled to observe the samples at the probe outlet and ensure that clean and blood-free samples were obtained. A cleaning cycle consisted of infusing 300 nL of perfusate into the tissue at 3 nL/s and sampling it back at the outlet. At the end of the session, the probe was removed, and the samples were stored in the outlet storage capillary at −20 °C, thus preserving their temporal information until analysis. The mouse was sutured and then allowed to recover. Blood glucose measurements were carried out by collecting 0.6 µL of blood from the tail vein on FreeStyle Precision test strips, with a glucometer FreeStyle Precision Neo (Abbott).

### 2.5. NanoESI-FTMS experiments

D-(+)-Glucose (G8270, CAS 50-99-7, Merck) in Perfusion Fluid CNS was used to prepare nanoESI-FTMS calibration samples. The additive was prepared by dissolving D-Glucose-^13^C6 (389374, CAS 110187-42-3, Merck) as internal standard (IS) in 67% deionized water and 33% acetonitrile (ACN, 900667, CAS 75-05-8, Merck). The LTQ Orbitrap ELITE ETD (Thermo Scientific) was equipped with the nanoSpray Flex Ion source (Thermo Scientific) and stainless steel nanoemitters PSSE-3 (30 mm, OD 150 µm, ID 30 µm, PepSep) under 2.6 kV spray voltage. An ABIRD system (ESI Source Solutions) blew filtered air over the MS inlet. FT-MS spectra were recorded in the reduced profile mode at a resolution set to 30000. 1 microscan was used with a maximum injection time of 1000 ms and the AGC was set to 5E5. Glucose and IS were detected in Selected Ion Monitoring (SIM) mode using a 20 *m/z* wide window centered on 206 *m/z*. Ions were extracted at 203.05 *m/z* and 209.07 *m/z* respectively, with a mass extraction window of 100 ppm. In a first step, the identities of the glucose and the IS were confirmed by accurate mass measurements, fragmentation by Collision Induced Dissociation (CID, respectively 28 eV and 35 eV) and assignment of the main fragment ions. The sample capillary (250/360 µm ID/OD) was interfaced to the emitter though a custom zero-dead-volume transparent connector. A PTFE drain around the emitter prevented PFD accumulation^44^. Flow rate was driven with a NeMESYS low-pressure syringe pump, with a 10 µL glass syringe (1701N, Hamilton) at 120 nL/min, through LMT-55 tubing (97618-09, VWR). Data analysis was performed on Xcalibur (Thermo Scientific) and MATLAB R2020b (MathWorks).

## 3. RESULTS

### 3.1. Development of the DoD approach

The brain is a dense tissue, in which the extracellular space represents only 20% of the volume and where cells are spaced by 10 to 100 nm^49^. This limits direct sampling of ECF to slow flow rates in the 1 nL/min range^51^. Therefore, microdialysis and push-pull sampling systems collect analytes by diffusion from the ECF to a dialysate. The recovered concentration of an analyte in the dialysate is reported relative to the true tissue concentration, by the recovery fraction η. With continuous sampling, when analytes get captured faster than the rate at which new molecules reach the sampling location by diffusion, a depletion layer is created in the tissue. This happens in microdialysis and push-pull probes operating continuously. Consequently, when sampling starts, η is initially at its local maximum and enters a transient period, until it reaches a steady state value, η_ss_, once the depletion layer is stabilized. Long transient durations lead to long experiments before exploitable samples are obtained, with η_ss_ levels that do not represent the true tissue concentration, but only a fraction of it. This prevents absolute quantification of the analytes in the tissue directly from the samples. Moreover, this means that sampling history influences the instantaneous sampling. Both η and the transient duration depend on the characteristics of the sampling system, on its operation and on the physiological interactions of the analyte with the tissue (molecule-specific diffusion, exchange and clearance processes in the tissue)^52–55^. Should any of these parameters change, a new equilibrium would need to be reached. Despite experimental (e.g. zero-net-flux) and computational methods to estimate η_ss_ and infer the real tissue concentration^54,56,57^, quantification remains tedious to perform and requires empirical parameters. The DoD approach was developed to solve the limitations above. It is a novel punctual push-pull sampling paradigm, complementary to continuous microdialysis and push-pull approaches. It proposes a discrete sampling sequence, implemented in a dedicated microprobe. DoD perfuses a controlled volume of perfusate, which collects analytes within an accurate diffusion window, before being sampled back as a droplet. This aims to provide a sample with high η and closely mirroring the tissue concentration at a defined time point, for quantitative molecular studies. Repeated samplings can be achieved by introducing an equilibration period between punctual sequences, to prevent creating a stable depletion layer causing a transient regime. This absence of transient regime makes acute sampling experiments possible, in less than an hour, and allows collecting independent droplet samples.

### 3.2. Fluidic probe for DoD

A dedicated microfluidic probe was developed to enable application of DoD in the brain of mice, as displayed in Figure 1. Inspired by previous work^25^, it is made of two flexible polymeric layers. The fabrication details are reported in Supporting Information (Figure S1). The polyimide layer forms the base that provides fluidic access to the tissue at the tip and contains the electrical functions of the probe (droplet-sensing electrodes and stimulation electrode), whereas the SU-8 layer defines the 40 µm high fluidic channels and the droplet formation geometry. A 40 µm wide perfusion channel enables delivery of perfusate to the tissue to collect molecules from the brain extracellular space by diffusion. The perfusate loaded with molecules is then aspirated from the tissue at the sampling aperture. It forms a droplet within the continuous PFD phase at the T-junction, where the 80 µm wide inlet and outlet channels filled with inert PFD intersect. PFD from the inlet then moves the droplet to the outlet, where the 20 µm wide, integrated platinum droplet-sensing electrodes detect it. All fluidic channels are interfaced to fused silica capillaries and the droplets reaching the outlet are transferred to a storage capillary, preserving their generation order until analysis.

**Figure 1.**
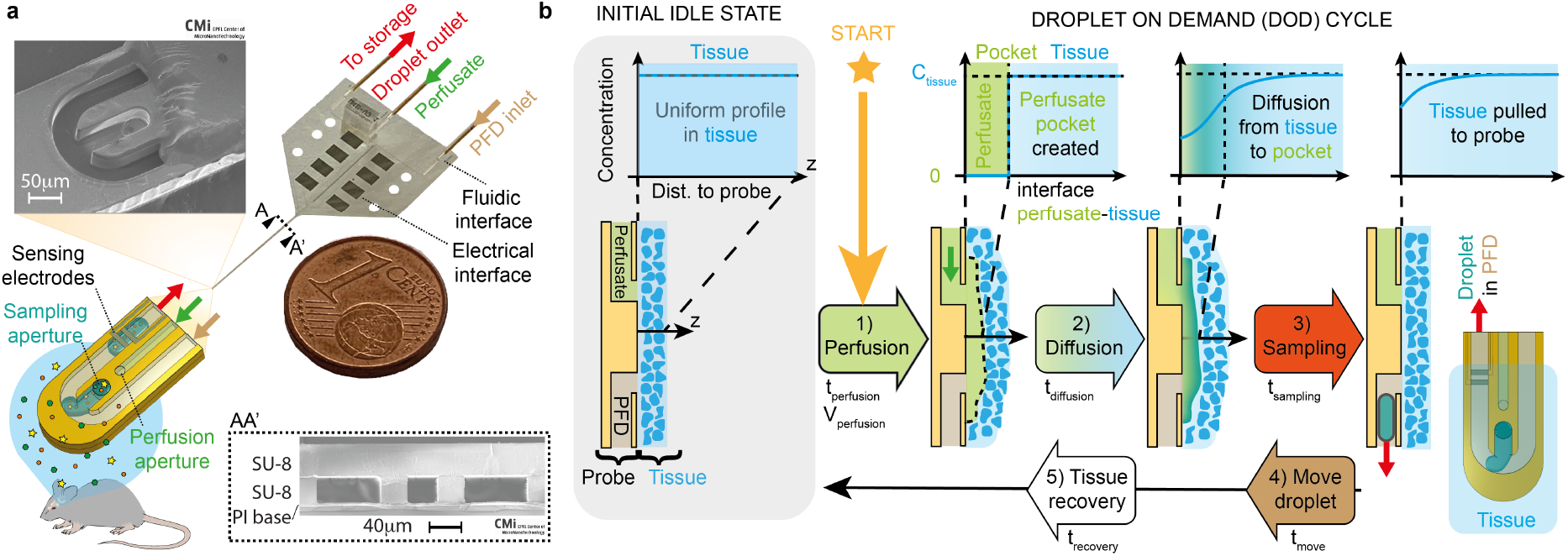
Implementation of the Droplet on Demand (DoD) method. **a** The flexible polyimide/SU-8 probe comprises three microfluidic channels with apertures at the tip of the needle inserted in the brain. They deliver perfusate and collect samples loaded with molecules from the extracellular space, as droplets separated by PFD. Electrodes detect them on the way to the storage capillary at the outlet. Scanning electron microscope pictures show the T-junction forming droplets at the tip and the cross-section of the needle, with the polyimide (PI) base embedding the electrodes, and the fluidic channels in SU-8. **b** Concept of the DoD approach implemented at the tip of the probe. Only the perfusion and outlet lines are represented. The tissue and the probe are initially considered idle, at rest, leaving the tissue unperturbed, with a uniform concentration profile of an analyte. 1) A punctual DoD sequence starts on demand with local infusion of perfusate, creating a pocket between the probe and the tissue. 2) The molecules diffuse from the tissue to the pocket. 3) The pocket is sampled and forms of a droplet within the PFD. 4) PFD moves it to the storage capillary. 5) To sample repeatedly, a recovery step is added to the punctual sequence (steps 1 to 4) for molecules to equilibrate their concentration in the tissue until the next cycle. This defines a DoD cycle.

### 3.3. Punctual DoD sequence and repetitive DoD cycles

The concept of the DoD approach is illustrated in Figure 1b as implemented at the tip of the probe, with the perfusion and sampling channels respectively filled with the non-miscible perfusate and PFD phases. The concentration of the analyte is considered spatially homogenous in the ECF, with no exchanges within the tissue. After inserting the probe in the tissue, the probe is idle, in contact with the tissue. A punctual DoD sequence is started on demand, by perfusing the tissue with a controlled volume of perfusate Vperfusion, over a duration tperfusion. Due to the density of the tissue, the perfusate forms a pocket of liquid at the probe-tissue interface, pushing the tissue away from the probe. The perfusate being free of analytes, a steep gradient exists that causes diffusion from the ECF to the pocket of perfusate. At the end of the diffusion time, defined as tdiffusion, the pocket is aspirated inside the probe during the sampling step, for a duration tsampling, and it forms a droplet within PFD, that is later moved to the outlet over tmove.

A punctual DoD sequence outputs one unique sample with a concentration corresponding to the average concentration of the pocket at the time of sampling, representing the tissue concentration over the window tdiffusion. At the punctual sequence level, a recovery fraction η close to 1 is aimed by adjusting Vperfusion and tdiffusion. By adding a recovery step of duration trecovery to the punctual DoD sequence, with all flows stopped, repetitive sampling can be performed, as DoD cycles. This lets the tissue equilibrate its concentration over an equilibration time tequilibration, defined in equation (1). It ensures that the next cycle will start with initial conditions nearly identical to the initial conditions of the previous one. This prevents the formation of a stable depletion layer in the tissue, that would result in a transient regime with a decay of η across subsequent samples. The temporal resolution is equal to tcycle, defined in equation (2).

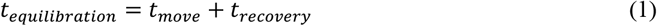

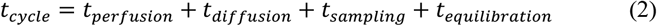

### 3.4. DoD simulations

The effects of tdiffusion and tequilibration over the recovery fraction of samples collected with DoD were studied in a passive tissue, with a finite element analysis model. It considered the tip of the needle at the center of a cube of 0.6% agarose containing a single analyte (glucose), and with a parallelepipedal 30 nL pocket of perfusate between them, covering the needle, as displayed in Figure S2a. Initially, the spatial concentration in the perfusate and in the agarose were homogeneously set to 0 µM and 1 µM of glucose respectively, with mass transport occurring by diffusion only. The spatial concentration in the system was simulated over time. Figure 2a reports the evolution of the average glucose concentration in the pocket over a single diffusion step, against tdiffusion, for pocket thicknesses from 20 to 160 µm. As expected, the simulations confirm that η increases with tdiffusion, even if longer tdiffusion is required to reach a η of 1. They also show that the increase rate of η reduces as η approaches 1. This mitigates the utility of increasing the diffusion time much more, especially for repeated sampling, if temporal resolution is important. In addition, the thinner the pocket of perfusate, the faster η increases, thus suggesting using small perfusion volumes. In real conditions, the effective thickness of the pocket of perfusate is uncontrolled and could influence the absolute results, but not the general evolution of η with respect to the diffusion time.

**Figure 2.**
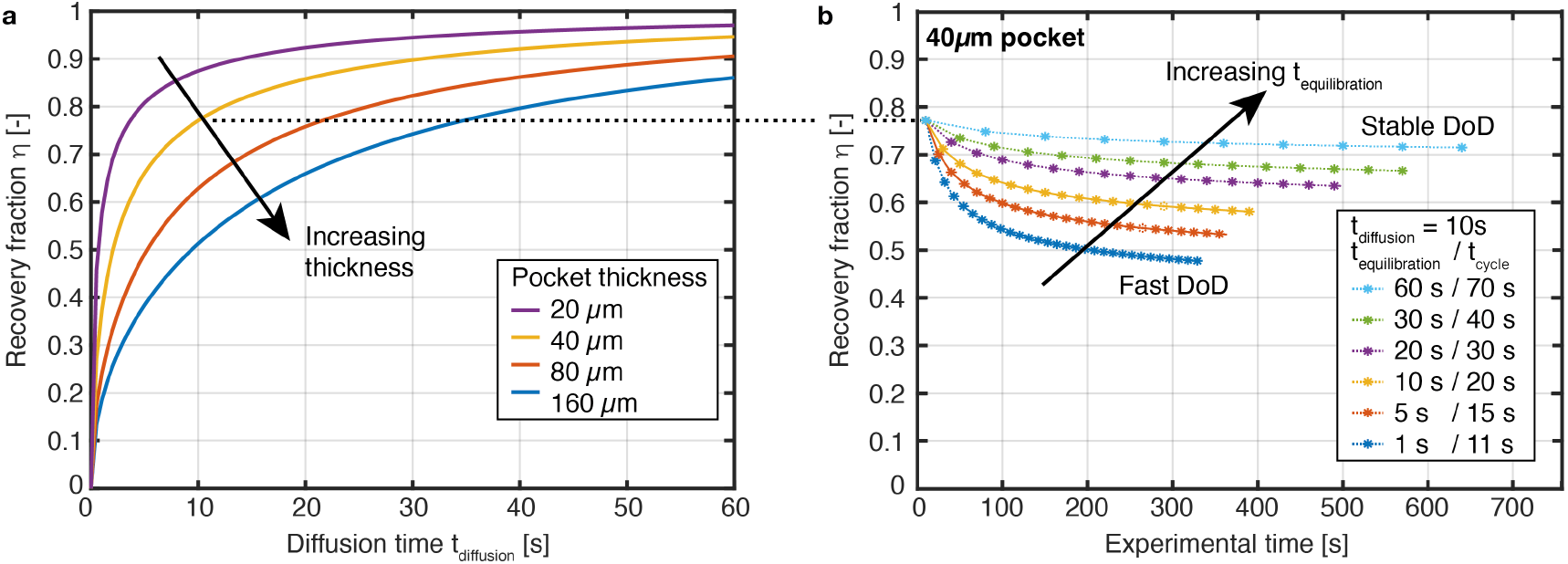
Simulations of DoD considering diffusion of glucose from an agarose brain phantom to a pocket of perfusate covering the needle of the probe. **a** Simulated punctual DoD sequence: evolution of the recovery fraction in the pocket during the diffusion step, as function of tdiffusion, for different pocket thicknesses. **b** Simulated DoD cycles applied repetitively and implemented in three steps: a diffusion step over 10 s of tdiffusion, instantaneous sampling, and variable equilibration step over 1 to 60 s of tequilibration between two samplings. The simulated recovery fraction of the collected samples is reported by stars connected by dashed lines to guide the eyes.

Repeated DoD cycles were simulated to study the effect of the equilibration time, by monitoring the evolution of η across samples. Based on the model above for punctual sampling, cycles were simulated repeatedly, in three steps: diffusion of molecules from the medium to the pocket of perfusate over tdiffusion, instantaneous sampling of the pocket and equilibration in the medium only, over tequilibration, until the next cycle. Using this model, the average η in the pocket at the sampling points is reported by stars in Figure 2b, for cycles that use a diffusion time of 10 s, an equilibration time from 1 to 60 s, and a 40 µm pocket of perfusate. All conditions provide a first sample as shown in the first diffusion step in Figure 2a. With short equilibration times of 1 s (fast DoD), η decreases across subsequent samples, thus illustrating of a short transient regime, similar to the one expected for continuous sampling. For 60 s of equilibration (stable DoD), the transient regime is significantly reduced, and η is maintained at a high level throughout the sampling process. In these conditions, repeated sampling cycles are equivalent to punctual DoD sequences. The longer tcycle reduces the temporal resolution, however this might not be critical unless fast variations are being tracked, depending on the study.

Overall, this confirms that increasing the diffusion and equilibration times enables sampling at high and steady η. This model assumes instantaneous pocket creation and sampling, but practically, diffusion into the perfusate would occur from the beginning of the perfusion to the end of the sampling. It is therefore adequate to study the concepts of DoD, but strict comparison to experimental data should be avoided.

### 3.5. DoD sampling *in vitro*

*In vitro* experiments verified the implementation of DoD in the probe, by sampling salts from an agarose gel brain phantom soaked in 150 mM NaCl, as in Figure 3a. The tip of the probe was inserted at the center of the gel to emulate sampling in the brain. It reproduces the hydraulic conditions in the brain better than fluid in a beaker, however it is less dense and does not integrate the exchanges and metabolism in the brain. Nonetheless, this is a valid model^22,50,58^ to confirm the features of the DoD. The inlet and perfusion lines were driven by syringe-pumps and respectively filled with PFD and an aqueous solution of 300 mM sucrose, as perfusate with low conductivity. Pressure control combined with a flow sensor at the outlet, provided a reactive fluidic system and limited the negative pressure applied to the gel, thus preventing direct sampling by ultrafiltration. A DoD cycle started by infusing perfusate at 5 nL/s to create a 30 nL pocket of perfusate in the gel. The diffusion time was varied from 1 to 60 s and the pocket was aspirated over 10 s at −3 nL/s by pulling the outlet to −150 mbar, as in Figure 3b. After aspiration, the droplet was moved to the outlet by circulating PFD within the probe at 3 nL/s, for 40 s. All flows were then stopped, and a recovery time of 20 s was observed, resulting in an equilibration time of 60 s before the next cycle. An example of actuation sequence is reported in Figure S3, with DoD parameters in Table S1. While sampling, the electrodes in the outlet channel measured the amplitude of the current across the droplets as in Figure 3c, which mirrors the conductivity, thus the recovered NaCl concentration.

**Figure 3.**
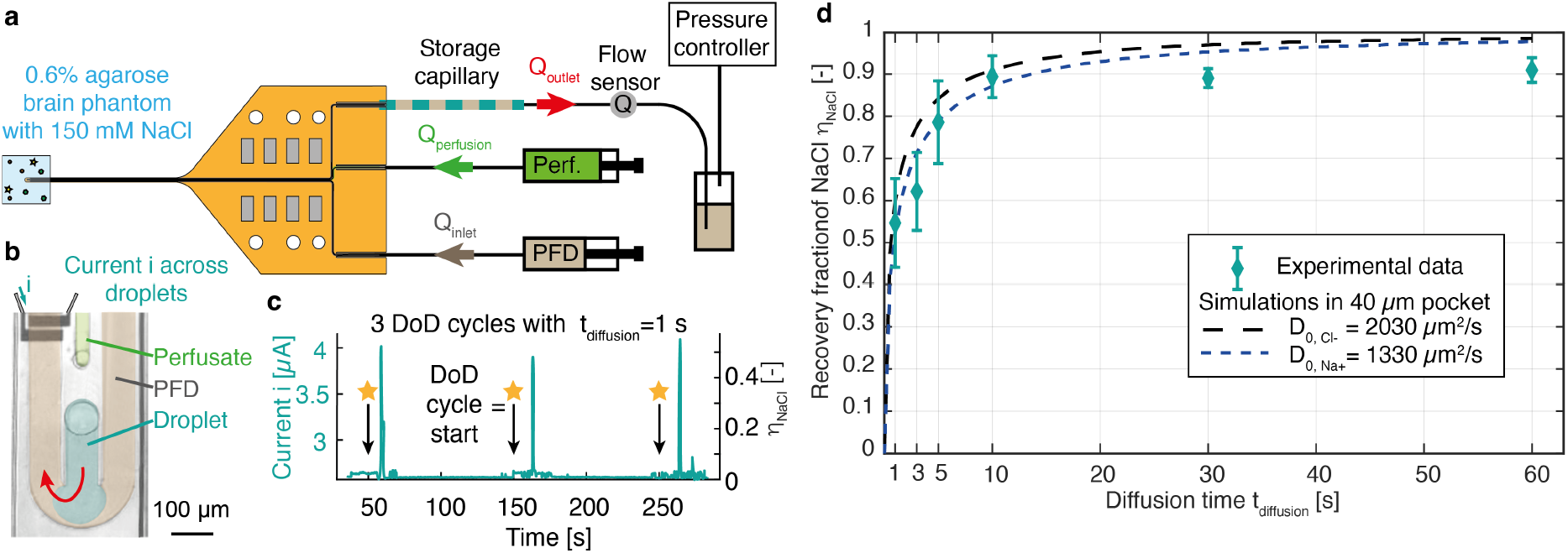
DoD sampling experiments in vitro. **a** Fluidic setup for sampling of NaCl from an agarose brain phantom. A syringe pump delivers perfusate into the gel, whereas the droplets are sampled by applying a negative pressure at the outlet. A second syringe pump drives the PFD moving the droplets to the outlet storage capillary. The flow sensor allows matching the inlet and outlet flow rates. **b** False-color image of the T-junction during a sampling step, with the aqueous phase aspirated within the PFD phase. The current is measured across the droplets by the electrodes in the channel. **c** The current amplitude across 3 subsequent droplets. The baseline current corresponds to PFD, whereas droplets generate a current pulse, that translates the recovery fraction of NaCl η_NaCl_. **d** Characterization of the effect of the diffusion time over η_NaCl_. The trend of the experimental data is compared to simulations considering the diffusion of Na+ or Cl^-^ into a 40 µm thick pocket of perfusate.

Figure 3d reports the recovery fraction of NaCl over the diffusion time and shows that the collected concentration increases with tdiffusion, until 10 s. The variability likely comes from the high sensitivity of the measured concentration to the diffusion time, since a strong concentration gradient exists between the pocket and the medium, and from the non-instantaneous sampling. For longer diffusion times, the recovery fraction reaches a plateau at 0.89±0.04, with reduced error bars, probably limited by the local dilution by the perfusate. Simulations with Na^+^ and Cl^-^ as the analytes, with a 40 µm thick pocket are added in Figure 3d. The real thickness of the pocket is unknown; however, it is expected to be within tens of microns, up to the thickness of the probe. This uncertainty prevents quantitative comparison, but the experimental data appears to follow a trend consistent with simulations. Moreover, no transient regime appeared in any of the DoD sampling conditions used (Figure S4a), even when the equilibration time was reduced to 6 s. This could be due to the thin pocket of perfusate and to the sampled volumes being small enough, so that the quantity of removed analyte was negligible, but also to the high diffusion coefficients of Na^+^ and Cl^-^ in water, 1330 and 2030 µm^2^/s, respectively. These results were verified to be specific to DoD, by using the probe in continuous sampling mode (Figure S4b). Opposite to DoD results, brief transient regimes were observed, together with lower η_ss_, dependent on the flow rate. These experiments verified that the high η and the absence of transient regime are inherent to DoD and thus validated the implementation of the probe *in vitro*.

### 3.6. DoD sampling *in vivo*

To demonstrate the potential of DoD in a complex *in vivo* setting, DoD was applied in the striatum of mice anesthetized with ketamine/xylazine. The setup and methods were similar to the ones of *in vitro* experiments. The inlet and outlet lines were primed with PFD, whereas the perfusion line used CNS perfusion fluid. The probe was secured on a holder mounted on a stereotactic frame as in Figure 4a and inserted into the brain of the animal, with a small hole in its skull. During insertion, all lines dispensed fluid at 3 nL/s to avoid clogging the channels and the flows were stopped once in position. Two cleaning cycles were performed to remove the dispensed fluids and potential debris, until clean samples were seen at the outlet. The probe was then left idle for 10 minutes, for the tissue to equilibrate. DoD cycles were then applied, with a sufficiently long equilibration time to consider them as individual sequences. During sampling, the droplet-sensing electrodes tracked the droplets and verified successful operation (Figure S5). The flow sensor at the outlet enabled adjusting the sampling duration and pressure, to compensate for changes of hydraulic resistance during the experiments. This prevented perfusing PFD to the brain or sampling by ultrafiltration (Figure S6). At the end, the probe was removed, and the animal was sutured back and allowed to recover. The procedure did not induce any noticeable impairment.

**Figure 4.**
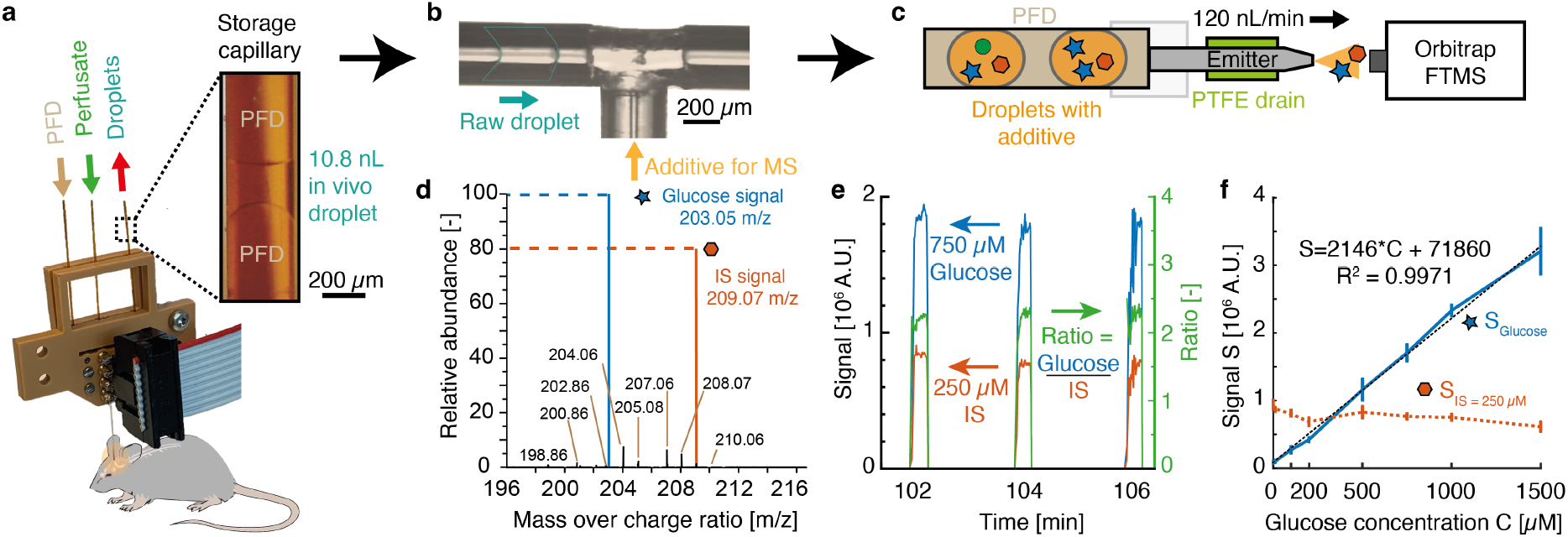
In vivo DoD experiments: from sampling to quantification of glucose in the droplets. **a** The probe sampled droplets from the mouse striatum, that were collected in the storage capillary at the outlet within PFD. **b** Picoinjection of the additive into the raw droplets at a custom-made junction. **c** Droplet analysis by direct infusion nanoESI-FTMS and electrospray of the analytes in front of the mass spectrometer. **d** Ion chromatogram showing the relative abundance of the detected species in in vivo samples. **e** 30 nL reference droplets of glucose and IS were used to calibrate the ratio between the two analytes. **f** Calibration of the glucose signal measured in reference samples with 250µm IS.

DoD cycles infusing 30 and 60 nL of perfusate enabled reliable droplet sampling (Movie S1), with characteristics in Table 1. Droplets smaller than 12 nL happened to split on the storage capillary walls, until collected by the next droplet, thus giving a temporal resolution of 1 to 2 t_cycle_. Functional treatment of the capillary prevented this, by reducing biofouling and the wetting. However, the effect fainted over time with *in vivo* samples, as illustrated by the contact angle between the droplet and the walls of the capillary in Figure 4a. Using a perfusion volume of 60 nL provided larger droplets, with a better sampling yield, defined as the rate of sequences producing a droplet successfully with respect to all the sequences applied. Although this increased droplet volume led to less droplets merging in the capillary, the larger perfusion volume could dilute them more, depending on the characteristics of the created pocket of perfusate.

**Table 1.** Characteristics of droplets sampled in vivo by DoD.

| Perfusion volume | Average droplet size | Standard deviation | Median droplet size | 1<sup>st</sup> quartile | 3<sup>rd</sup> quartile | Number of droplets | Sampling yield |
| --- | --- | --- | --- | --- | --- | --- | --- |
| [nL] | [nL] | [nL] | [nL] | [nL] | [nL] | [-] | [%] |
| 30 | 9.9 | 4.1 | 9.6 | 7.2 | 11.4 | 88 | 85 |
| 60 | 24.4 | 15.7 | 21.0 | 12.2 | 30.7 | 115 | 91 |

### 3.7. Quantification of glucose

NanoESI-FTMS was selected to analyze the content of the droplets, not only because of its multiplexing and absolute quantification capabilities, but also for its sensitivity, its compatibility with nanoliter-sized samples separated by perfluorodecalin, and its robustness to ion suppression^42,44^. Glucose is essential to the brain and presents metabolic dysfunctions in multiple neurodegenerative diseases^59^, such as upon aggregation of amyloid-β peptides^60^. Therefore, it proved a suitable metabolite to demonstrate the performances of DoD. Glucose could be detected by direct infusion of raw *in vivo* samples, but the complex sample matrix and the absence of separation method prevented direct quantification reliably. Therefore, an additive containing a glucose isotope,^13^C6H12O6, prepared in deionized water and ACN, was used as an internal standard (IS). After sampling, the additive was added to the raw droplets by picoinjection using the custom-made T-junction in Figure 4b, thus diluting 10 times the glucose and the salts inducing ion-suppression. The final IS concentration was set close to the final glucose concentration and the final ACN content to 30%. Droplets with additive were infused through the emitter under high voltage and sprayed into the mass spectrometer as Figure 4c. The signals of glucose and IS were monitored as [M+Na]^+^ adducts at 203.053 *m/z* and 209.073 *m/z* respectively, and their relative abundance was measured as in Figure 4d. The median ratio over the passage of droplets was evaluated (Figure 4e), and a calibration with reference droplet samples was acquired (Figure 4f). The signal of glucose depended linearly on the glucose concentration, whereas the signal of 250 µM IS did not vary within the final 100-1000 µM glucose range. Thus, the IS was a suitable reference for absolute quantification of glucose.

### 3.8. Glucose in brain samples

The effect of the diffusion time on the glucose concentration collected *in vivo* was studied. Figure 4 shows the method used to quantify glucose sampled in the brain of mice with DoD. Figure 5a reports the glucose concentration of the 10 first subsequently collected droplets of a sampling session with an animal, using a diffusion time of 10 s, with 30 nL of perfusate and parameters in Table S1. The insets show the nanoESI-FTMS signals obtained for glucose and 200 µM IS, and their ratio. No clear decay of the concentration, typical of a transient regime, was observed across the samples and the mean concentration of glucose was quantified at 2.73±0.35 mM. Reducing tdiffusion to 1 s reduced the concentration of recovered glucose, as reported in Figure 5b. Using 60 nL of Vperfusion provided a similar trend, with the mean glucose concentration reaching 3.48±0.17 mM after 60 s of diffusion, without any transient effect observed. For a diffusion time of 10 s, the concentration was slightly lower with Vperfusion of 60 nL than with 30 nL. This was not investigated further but could be due to the perfusion volume influencing the shape of the pocket, or to the insertion affecting the contact between the probe and the tissue, or to different glucose concentrations in the animals.

**Figure 5.**
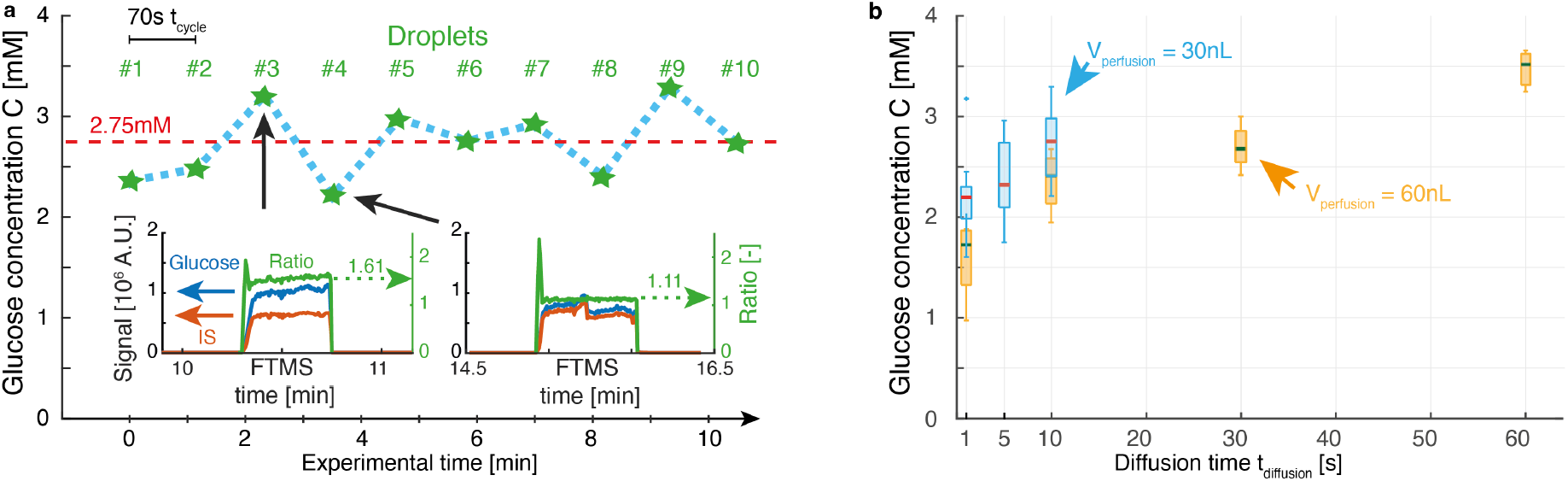
Glucose measured from droplets collected in vivo. **a** The concentration of glucose is measured in the first 10 subsequently sampled droplets, using a perfusion volume of 30 nL, a diffusion time of 10 s and an equilibration time of 30 s. The dashed line represents the median concentration of 2.75 mM. The insets show the signals of glucose and IS by nanoESI-FTMS, with their median ratio. **b** Glucose concentration measured in droplets sampled with perfusion volumes of 30 nL and 60 nL, as a function of the diffusion time used by the DoD sequence.

## 4. Discussion

A new sampling approach was inspired by the limitations of state-of-the-art continuous sampling methods. The proposed DoD sequence was described (Figure 1) and implemented in a microfabricated probe, then confirmed by simulations and samplings *in vitro*. The simulations in Figure 2 illustrate the effects of the thickness of the perfusate pocket and of the diffusion and equilibration times on the recovery fraction, for punctual and repeated DoD samplings. The *in vitro* samplings reported in Figure 3 confirmed the increasing recovery fraction with the diffusion time, predicted by simulations. An appropriate choice of perfusion volume, diffusion and equilibration times provided a high and steady recovery fraction across samples, equivalent to no transient regime. These characteristics are unique to the DoD method.

DoD was also applied *in vivo* and an analytical protocol with nanoESI-FTMS was developed. Herein, we demonstrated that DoD successfully enabled collecting samples from the brain over sampling windows of 60 to 120 minutes, depending on the duration of the anesthesia. 30 and 60 nL perfusion volumes resulted in respective sampling yields of 85% and 91%. Despite the brain surface did not show any backflow leaking out, only 33% and 40% of the perfusion volumes were sampled back respectively (Table 1). Due to the density of the brain, a negligible fraction of the perfusate was expected to penetrate the tissue and the perfusion rate was kept low to avoid backflow of perfusate^61^. Although we did not observe any leak, the missing volume of perfusate infused at the tip has likely flown back along the needle, towards the surface of the brain. This highlights the need to further investigate the formation of the pocket of perfusate in the tissue and optimize the perfusion volume. The appropriate parameters are influenced by the mass transport of the analytes. When sampling repeatedly, they directly determine the temporal resolution, given by the cycle time. Reducing the fluidic volumes could improve it, thus requiring a shorter diffusion time. However, not all analytical methods are suited for small samples and not all analytes require fast sampling^13^.

Quantification of glucose in the droplets collected *in vivo* showed that the DoD method is suited for sampling molecules in the brain (Figure 4). The effect of the diffusion time over the recovered concentration, predicted by simulations and observed *in vitro*, was confirmed *in vivo* (Figure 5). No transient regime was observed in any of the conditions and therefore the minimal required equilibration time was not investigated. Consequently, the first sample is already representative of the tissue concentration, and in these conditions, a reliable sample can be obtained within a sampling session as short as 20 minutes.

The glucose concentration in the brain was reported in the 0.3 to 3.3 mM range in the literature, measured with many methods^62^ and in different brain compartments in rats^63^. The values measured in this work (Figure 5) are close to the highest reported values, and higher than the concentrations of 350±20^64^ and 530 µM^65^ reported in the striatum of awake rats with microdialysis. Similar values are expected in the mouse striatum and the difference is likely explained by a hyperglycemia induced by the anesthesia, as verified by blood glucose measurements from the tail vein of an animal. In the blood, a glycemia of 8.1 mM was measured 5 minutes after anesthesia and it increased around 23.8±3.2 mM over the duration of the sampling session. These measurements suggest that blood glucose levels increased to at least 294% of the baseline euglycemia level. This is in agreement with the reported hyperglycemia level at 268% of the baseline 60 minutes after anesthesia with ketamine/xylazine, which reached up to 460% of the baseline in some cases^66^. This can explain the high concentration measured in the samples and suggests that the hyperglycemia induced in the bloodstream by the anesthesia is reflected in the striatum. Not knowing the exact tissue concentration in these conditions makes it difficult to estimate the true recovery fraction of glucose. However, based on the measured values and on the literature, it is probably high. Moreover, like observed *in vitro*, the effect of the diffusion time is clear, and no transient regime was observed, thus validating the implementation of DoD *in vivo*.

## 5. CONCLUSIONS

DoD was demonstrated as a new sampling paradigm through *in vitro* and *in vivo* experiments. As a proof of concept, DoD was applied to quantify the glucose in the striatum, by nanoESI-FTMS, performed on the retrieved droplets. The results confirmed that the hyperglycemia induced in the blood-stream by ketamine/xylazine is reflected in the striatum. Sampling in awake and freely moving animals should be possible as it would only require updating the probe and the animal license, that were intended for acute experiments in anesthetized animals.

DoD can be tailored to the needs of the *in vivo* study and by adequate choice of the parameters, samples that closely mirror the true tissue concentration can be obtained for absolute quantification. This should be optimized according to the target molecules. In addition to glucose, glutamate and acetylcholine were detected in DoD samples, and their quantification will be optimized for future studies. In parallel, the characteristics of the pocket of perfusate will also be investigated.

Among the fluidic sampling solutions, DoD offers sequential sampling, with similarities to the proposed, but, to the knowledge of the authors, never implemented digital microdialysis approaches^67,68^. It features on demand sampling at high recovery fraction, and outputs one sample per punctual sequence. Multiple samples with high and stable recovery fraction can be obtained by repetition, while letting the tissue equilibrate between samplings. The collected samples can later be used independently for statistics or to track an absolute variation over time. They can also be merged to provide a single, large, and highly concentrated sample.

Due to the absence of transient regime, the first sample is already representative of the ECF content, and subsequent samples are not influenced by the previous sampling history. This allows experiments as short as 20 minutes for a single sample and collection of multiple relevant samples with a temporal resolution of 1-2 minutes over longer sampling sessions. The exact performances will depend on the target analytes, but these features distinguish DoD from continuous sampling methods with a transient period, that use droplets to improve temporal resolution. Instead, DoD is a novel and complementary method that provides droplet samples mirroring the tissue concentration at each cycle, thus enabling absolute quantification while reducing the duration of experiments. Performing DoD and microdialysis in parallel could provide further insights on the potential of DoD and improve the calibration of micro-dialysis probes.

The current implementation could find direct applications in the study of volume transmission and hormone fluctuation events^13^, and the absence of transient regime could improve the study of pharmacokinetic transients following a bolus or a steady infusion^52^. Contrary to obtaining a high recovery fraction, reaching a higher temporal resolution requires reducing the cycle time. Reducing the fluidic volumes might enable to meet both criteria but suitable analytical methods should be available to process such smaller samples^69^.

The absence of transient could possibly enable quantifying analytes with slow accumulation dynamics and mass transport, that would otherwise be discarded during the transient period of continuous sampling methods and of which the recovery fraction could be too low for reliable measurement. The absence of membrane on the DoD probe could allow collection of large biomarkers^20^, such as extracellular vesicles^70^. Having a relatively low abundance, they would benefit from the high recovery fraction offered by DoD. These biomarkers are rich in analytes, namely microRNAs^71^, which could provide valuable insights on brain pathologies and guide efficient treatments. Fast access to samples mirroring the true tissue concentration could improve the diagnostic and monitoring of neurodegenerative diseases, which develop slowly, and induce molecular concentration changes and protein aggregation in the brain. Obviously, the DoD approach is applicable to other tissues.

## 6. SUPPORTING INFORMATION

Supporting Information: Fabrication process; DoD simulations; Flow actuation diagram; DoD sampling *in vitro* – concentration of subsequent DoD samples; DoD sampling *in vivo* – electrical droplet detection; DoD sampling *in vivo* – flow rate; DoD sampling *in vivo* – parameters of movie of 6 DoD cycles, Table of sampling parameters for experiments (PDF file).

Supporting Movie “6 DoD cycles *in vivo*” (MP4 file).

## 7. AUTHOR CONTRIBUTIONS

P.R. and A.B. proposed the study, supervised the project, and discussed the results with J.T. J.T. did the DoD simulations, fabricated the probes, conducted the *in vitro* and *in vivo* experiments, analyzed the data, and wrote the manuscript. S.N. performed the surgery for *in vivo* experiments. J.T., L.M. and D.O. developed the method for droplet analysis by nanoESI-FTMS. Funding for *in vivo* experiments was provided by H.L. The rest was funded by P.R. All authors commented on the manuscript.

## 8. ACKNOWLEDGMENT

The authors are thankful to the staff of the EPFL Center for MicroNanoTechnology (CMi) for their help optimizing the fabrication process and for the quality of the platform, whereas G. Petit-Pierre, S. Jiguet and C. Hibert are thanked for their precious advice on the use and fabrication of the probes. The authors thank the staff of the EPFL Center of PhenoGenomics for their support and the staff of the MSEAP for their precious help developing the analytical method. S.N. and J. Burtscher are thanked for their advice and work on preparing the *in vivo* license and experiments. The authors acknowledge EPFL Internal Fund and grant #591-579.

## REFERENCES

1. Zhang, Y., Jiang, N. & Yetisen, A. K. Brain Neurochemical Monitoring. Biosens. Bioelectron. 189, 113351 (2021).

2. Mercanzini, A., Dransart, A. & Pollo, C. Directional Deep Brain Stimulation. In Innovative Neuromodulation, 61–82 (2017).

3. Schwerdt, H. N. et al. Subcellular probes for neurochemical recording from multiple brain sites. Lab Chip 17, 1104–1115 (2017).

4. Shin, H. et al. 3D high-density microelectrode array with optical stimulation and drug delivery for investigating neural circuit dynamics. Nat. Commun. 12, 492 (2021).

5. Chae, U. et al. Bimodal neural probe for highly co-localized chemical and electrical monitoring of neural activities in vivo. Biosens. Bioelectron. 191, 113473 (2021).

6. Burton, A. et al. Wireless, battery-free, and fully implantable electrical neurostimulation in freely moving rodents. Microsystems Nanoeng. 7, 62 (2021).

7. Minev, I. R. et al. Electronic dura mater for long-term multimodal neural interfaces. Science 347, 159–163 (2015).

8. Canales, A. et al. Multifunctional fibers for simultaneous optical, electrical and chemical interrogation of neural circuits in vivo. Nat. Biotechnol. 33, 277–284 (2015).

9. VanDersarl, J. J., Mercanzini, A. & Renaud, P. Integration of 2D and 3D Thin Film Glassy Carbon Electrode Arrays for Electrochemical Dopamine Sensing in Flexible Neuroelectronic Implants. Adv. Funct. Mater. 25, 78–84 (2015).

10. Jeong, J. W. et al. Wireless Optofluidic Systems for Programmable In Vivo Pharmacology and Optogenetics. Cell 162, 662–674 (2015).

11. Vázquez-Guardado, A., Yang, Y., Bandodkar, A. J. & Rogers, J. A. Recent advances in neurotechnologies with broad potential for neuroscience research. Nat. Neurosci. 23, 1522–1536 (2020).

12. Ngernsutivorakul, T., White, T. S. & Kennedy, R. T. Microfabricated Probes for Studying Brain Chemistry: A Review. ChemPhysChem 19, 1128–1142 (2018).

13. Frank, J. A., Antonini, M.-J. & Anikeeva, P. Next-generation interfaces for studying neural function. Nat. Biotechnol. 37, 1013–1023 (2019).

14. Sim, J. Y., Haney, M. P., Park, S. Il, McCall, J. G. & Jeong, J.-W. Microfluidic neural probes: in vivo tools for advancing neuroscience. Lab Chip 17, 1406–1435 (2017).

15. Le Roux, L. G. & Schellingerhout, D. Molecular Neuroimaging: The Basics. Semin. Roentgenol. 49, 225–233 (2014).

16. Nasrallah, I. & Dubroff, J. An Overview of PET Neuroimaging. Semin. Nucl. Med. 43, 449–461 (2013).

17. Metz, S., Jiguet, S., Bertsch, A. & Renaud, P. Polyimide and SU-8 microfluidic devices manufactured by heat-depolymerizable sacrificial material technique. Lab Chip 4, 114 (2004).

18. Lee, W. H. et al. Microfabrication and in Vivo Performance of a Microdialysis Probe with Embedded Membrane. Anal. Chem. 88, 1230–1237 (2016).

19. Lee, W. H., Slaney, T. R., Hower, R. W. & Kennedy, R. T. Microfabricated sampling probes for in vivo monitoring of neurotransmitters. Anal. Chem. 85, 3828–3831 (2013).

20. Wu, G. et al. Wireless, battery-free push-pull microsystem for membrane-free neurochemical sampling in freely moving animals. Sci. Adv. 8, (2022).

21. Wang, M., Roman, G. T., Schultz, K., Jennings, C. & Kennedy, R. T. Improved Temporal Resolution for in Vivo Microdialysis by Using Segmented Flow. Anal. Chem. 80, 5607–5615 (2008).

22. Raman, R. et al. Platform for micro-invasive membrane-free biochemical sampling of brain interstitial fluid. Sci. Adv. 6, eabb0657 (2020).

23. van den Brink, F. T. G. et al. A miniaturized push–pull-perfusion probe for few-second sampling of neurotransmitters in the mouse brain. Lab Chip 19, 1332–1343 (2019).

24. Feng, S., Clement, S., Zhu, Y., Goldys, E. M. & Inglis, D. W. Microfabricated needle for hydrogen peroxide detection. RSC Adv. 9, 18176–18181 (2019).

25. Petit-Pierre, G., Bertsch, A. & Renaud, P. Neural probe combining microelectrodes and a droplet-based microdialysis collection system for high temporal resolution sampling. Lab Chip 16, 917–924 (2016).

26. Petit-Pierre, G. et al. In vivo neurochemical measurements in cerebral tissues using a droplet-based monitoring system. Nat. Commun. 8, 1239 (2017).

27. Basova, E. Y. & Foret, F. Droplet microfluidics in (bio)chemical analysis. Analyst 140, 22–38 (2015).

28. Shang, L., Cheng, Y. & Zhao, Y. Emerging Droplet Microfluidics. Chem. Rev. 117, 7964–8040 (2017).

29. Rhee, M. et al. Pressure stabilizer for reproducible picoinjection in droplet microfluidic systems. Lab Chip 14, 4533–4539 (2014).

30. Niu, X., Pereira, F., Edel, J. B. & de Mello, A. J. Droplet-Interfaced Microchip and Capillary Electrophoretic Separations. Anal. Chem. 85, 8654–8660 (2013).

31. Sun, X., Tang, K., Smith, R. D. & Kelly, R. T. Controlled dispensing and mixing of pico-to nanoliter volumes using on-demand droplet-based microfluidics. Microfluid. Nanofluidics 15, 117–126 (2013).

32. Piendl, S. K. et al. Integration of segmented microflow chemistry and online HPLC/MS analysis on a microfluidic chip system enabling enantioselective analyses at the nanoliter scale. Lab Chip 21, 2614–2624 (2021).

33. Liu, W. & Zhu, Y. “Development and application of analytical detection techniques for droplet-based microfluidics”-A review. Anal. Chim. Acta 1113, 66–84 (2020).

34. Zhu, Y. & Fang, Q. Analytical detection techniques for droplet microfluidics—A review. Anal. Chim. Acta 787, 24–35 (2013).

35. Feng, S., Shirani, E. & Inglis, D. W. Droplets for sampling and transport of chemical signals in biosensing: A review. Biosensors 9, 1–14 (2019).

36. Ha, N. S. et al. Faster, better, and cheaper: harnessing microfluidics and mass spectrometry for biotechnology. RSC Chem. Biol. 2, 1331–1351 (2021).

37. Leroy, A., Teixidor, J., Bertsch, A. & Renaud, P. In-flow electrochemical detection of chemicals in droplets with pyrolysed photoresist electrodes: application as a module for quantification of microsampled dopamine. Lab Chip 21, 3328–3337 (2021).

38. Suea-Ngam, A., Rattanarat, P., Chailapakul, O. & Srisa-Art, M. Electrochemical droplet-based microfluidics using chip-based carbon paste electrodes for high-throughput analysis in pharmaceutical applications. Anal. Chim. Acta 883, 45–54 (2015).

39. Slaney, T. R. et al. Push–Pull Perfusion Sampling with Segmented Flow for High Temporal and Spatial Resolution in Vivo Chemical Monitoring. Anal. Chem. 83, 5207–5213 (2011).

40. Nightingale, A. M. et al. Monitoring biomolecule concentrations in tissue using a wearable droplet microfluidic-based sensor. Nat. Commun. 10, 1–12 (2019).

41. Wang, M., Roman, G. T., Perry, M. L. & Kennedy, R. T. Microfluidic chip for high efficiency electrophoretic analysis of segmented flow from a microdialysis probe and in vivo chemical monitoring. Anal. Chem. 81, 9072–9078 (2009).

42. Ngernsutivorakul, T., Steyer, D. J., Valenta, A. C. & Kennedy, R. T. In Vivo Chemical Monitoring at High Spatiotemporal Resolution Using Microfabricated Sampling Probes and Droplet-Based Microfluidics Coupled to Mass Spectrometry. Anal. Chem. 90, 10943–10950 (2018).

43. Peretzki, A. J. et al. How electrospray potentials can disrupt droplet microfluidics and how to prevent this. Lab Chip 20, 4456–4465 (2020).

44. Beulig, R. J. et al. A droplet-chip/mass spectrometry approach to study organic synthesis at nanoliter scale. Lab Chip 17, 1996–2002 (2017).

45. Song, P., Hershey, N. D., Mabrouk, O. S., Slaney, T. R. & Kennedy, R. T. Mass Spectrometry “Sensor” for in Vivo Acetylcholine Monitoring. Anal. Chem. 84, 4659–4664 (2012).

46. Persike, M., Zimmermann, M., Klein, J. & Karas, M. Quantitative Determination of Acetylcholine and Choline in Microdialysis Samples by MALDI-TOF MS. Anal. Chem. 82, 922–929 (2010).

47. Bell, S. E. et al. Droplet Microfluidics with MALDI-MS Detection: The Effects of Oil Phases in GABA Analysis. ACS Meas. Sci. Au 1, 147–156 (2021).

48. Hartner, N. T. et al. Coupling Droplet Microfluidics with Ion Mobility Spectrometry for Monitoring Chemical Conversions at Nanoliter Scale. Anal. Chem. 93, 13615–13623 (2021).

49. Syková, E. & Nicholson, C. Diffusion in Brain Extracellular Space. Physiol. Rev. 88, 1277–1340 (2008).

50. Chen, Z.-J. et al. A realistic brain tissue phantom for intraparenchymal infusion studies. J. Neurosurg. 101, 314–322 (2004).

51. Kennedy, R. T., Thompson, J. E. & Vickroy, T. W. In vivo monitoring of amino acids by direct sampling of brain extracellular fluid at ultralow flow rates and capillary electrophoresis. J. Neurosci. Methods 114, 39–49 (2002).

52. Bungay, P. M., Dedrick, R. L., Fox, E. & Balis, F. M. Probe Calibration in Transient Microdialysis In Vivo. Pharm. Res. 18, 361–366 (2001).

53. Morrison, P. F. et al. Quantitative microdialysis. In Techniques in the Behavioral and Neural Sciences vol. 7 47–80 (1991).

54. Bungay, P. M., Morrison, P. F., Dedrick, R. L., Chefer, V. I. & Zapata, A. Chapter 2.2 Principles of quantitative microdialysis. In Handbook of Behavioral Neuroscience vol. 16 131–167 (2006).

55. Bungay, P. M., Morrison, P. F. & Dedrick, R. L. Steady-state theory for quantitative microdialysis of solutes and water in vivo and in vitro. Life Sci. 46, 105–119 (1990).

56. Stanley, M., Walt, H. & Joseph, B. J. In Vivo Calibration of Microdialysis Probes for Exogenous Compounds. Anal. Chem. 64, 577–583 (1992).

57. Bungay, P. M., Sumbria, R. K. & Bickel, U. Unifying the mathematical modeling of in vivo and in vitro microdialysis. J. Pharm. Biomed. Anal. 55, 54–63 (2011).

58. Pomfret, R., Miranpuri, G. & Sillay, K. The Substitute Brain and the Potential of the Gel Model. Ann. Neurosci. 20, 118–122 (2013).

59. Han, R., Liang, J. & Zhou, B. Glucose Metabolic Dysfunction in Neurodegenerative Diseases— New Mechanistic Insights and the Potential of Hypoxia as a Prospective Therapy Targeting Metabolic Reprogramming. Int. J. Mol. Sci. 22, 5887 (2021).

60. Allaman, I. et al. Amyloid-Aggregates Cause Alterations of Astrocytic Metabolic Phenotype: Impact on Neuronal Viability. J. Neurosci. 30, 3326–3338 (2010).

61. Morrison, P. F., Chen, M. Y., Chadwick, R. S., Lonser, R. R. & Oldfield, E. H. Focal delivery during direct infusion to brain: role of flow rate, catheter diameter, and tissue mechanics. Am. J. Physiol. Integr. Comp. Physiol. 277, R1218–R1229 (1999).

62. Routh, V. H. Glucose-sensing neurons: Are they physiologically relevant? Physiol. Behav. 76, 403–413 (2002).

63. McNay, E. C. & Gold, P. E. Extracellular glucose concentrations in the rat hippocampus measured by zero-net-flux: Effects of microdialysis flow rate, strain, and age. J. Neurochem. 72, 785–790 (1999).

64. Fray, A. E., Boutelle, M. & Fillenz, M. Extracellular glucose turnover in the striatum of unanaesthetized rats measured by quantitative microdialysis. J. Physiol. 504, 721–726 (1997).

65. Valenta, A. C., D’Amico, C. I., Dugan, C. E., Grinias, J. P. & Kennedy, R. T. A microfluidic chip for on-line derivatization and application to in vivo neurochemical monitoring. Analyst 146, 825–834 (2021).

66. Brown, E. T., Umino, Y., Loi, T., Solessio, E. & Barlow, R. Anesthesia can cause sustained hyperglycemia in C57/BL6J mice. Vis. Neurosci. 22, 615–618 (2005).

67. Chen, C. & Drew, K. L. Droplet-based microdialysis—Concept, theory, and design considerations. J. Chromatogr. A 1209, 29–36 (2008).

68. Chen, C.-F. Dimensional Analysis and Constitutive Equations of Quantitative Microdialysis. Insights Biomed. Res. 1, 5–11 (2017).

69. Hammarlund-Udenaes, M. Microdialysis for pharmacokinetic and pharmacodynamic studies with focus on the CNS. In Compendium of In Vivo Monitoring in Real-Time Molecular Neuroscience 47–69 (2017).

70. Stangler, L. A. et al. Microdialysis and microperfusion electrodes in neurologic disease monitoring. Fluids Barriers CNS 18, 1–14 (2021).

71. Bache, S. et al. Detection and quantification of microRNA in cerebral microdialysate. J. Transl. Med. 13, 149 (2015).

