## Supplementary Information for "On demand nanoliter sampling probe for collection of brain fluid"

### FABRICATION PROCESS

The dedicated DoD probe was fabricated at the EPFL Center of MicroNanoTechnology CMi and is illustrated in Figure S1a. The fabrication starts on a silicon wafer with a sputtered film of 100 nm W:Ti (10% wt) and 400 nm Al (Pfeiffer SPIDER600) as adhesive and sacrificial layers respectively (1). A 3.5 µm base layer of polyimide layer (PI 2611, DuPont) is spun coated and patterned with slanted walls by photolithography and dry etching process (STS Multiplex ICP). A film of 75 nm Ti, 350 nm Pt and 75 nm Ti is sputtered (Pfeiffer SPIDER600) and patterned by photolithography and ion beam etching (Veeco Nexus IBE350) to produce droplet-sensing electrodes (2). Electrodes are then passivated under a 1 µm secondary polyimide layer (PI 2610, DuPont), patterned similarly to the first layer (3). After 30 s oxygen plasma activation of the polyimide layer at 600 W (Tepla Gigabatch), SU-8 is spun coated and microfluidic channels (SU-8 3025, MicroChem or GM1070PI, Gersteltec) are defined by photolithography. The timing of SU-8 development is critical to ensure full development and avoid leakages between the channels. It is performed in 2 beakers of PGMEA, 1 minute in each one and with fresh PGMEA dispensed on the wafer surface every 30 s (4). To close the channels, a SU-8 cover layer is spun coated on a flexible and clear PET film (Levsurf LS100, Kimoto) on a secondary wafer. After 30 s oxygen plasma activation of the channels at 600 W, the cover layer is laminated to the SU-8 channels at 42°C (36 kg, 0.2 m/min, Bungard RLM419). Exposure of the SU-8 cover layer is performed through the PET film on a mask aligner (Süss MA6Gen3), through a photomask. The post-exposure bake is performed on a hotplate with the wafer stack upside-down, before peeling off the PET film at 65°C and developing the cover in PGMEA (5). The channels are then cut open with a laser (OPTEC LSV3, EXCIMER Laser 193 nm) prior to proceeding to the anodic release of the implants in a 2 M NaCl bath, by applying 0.7 V to the wafer with respect to a platinum counter electrode, 1 cm away (6). Implants are then cleaned in 1% HF for 1 minute to remove the outer Ti layer on the electrodes and expose the platinum. A scanning electron microscope (SEM) picture of the cross-section of the needle is reported in Figure S1b, showing 3 parallel channels and their dimensions. The fluidic interface is achieved by fitting the end of the microchannels inside fused silica capillaries (TSP-250350, Molex). UV sensitive epoxy glue (60-7170, EPOXIES ETC.) is applied to ensure sealing with no dead volume and no air traps at the channel-capillary interface as illustrated in Figure S1c. Curing is performed trough an optical fiber with a UV source (UVP SpotCure).

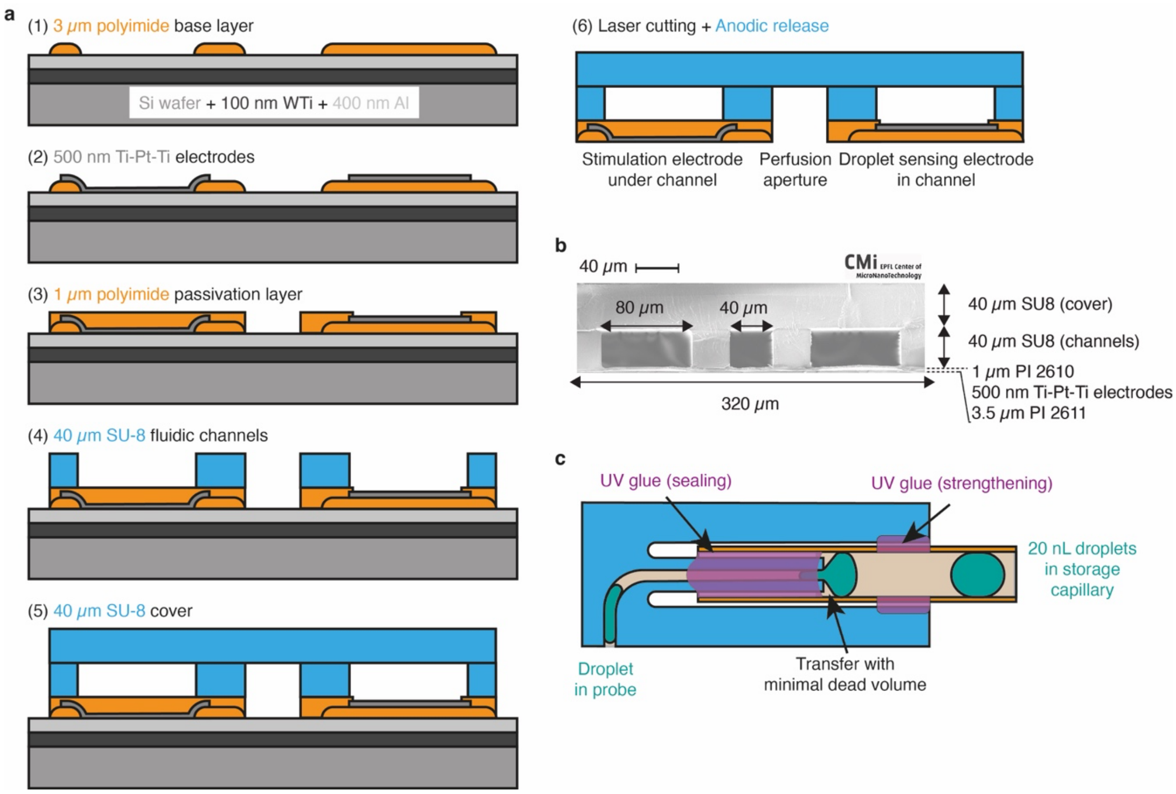


***Figure S1*** *Probe fabrication and interfacing.* ***a*** *Fabrication process: (1) patterning of the base polyimide layer, (2) patterning of the electrodes for external contact and internal contact, (3) patterning of the second polyimide layer, (4) patterning of the fluidic channels, (5) patterning of the cover to close the channels, (6) laser cutting to open the end of the channels and anodic release in NaCl bath.* ***b*** *SEM picture of a cross-section of the needle, showing the outlet, perfusion and inlet channels.* ***c*** *Schematic of the fluidic interface to transfer the droplets in the probe to the storage capillary and sealing with UV-curable epoxy.*

### DOD SIMULATIONS

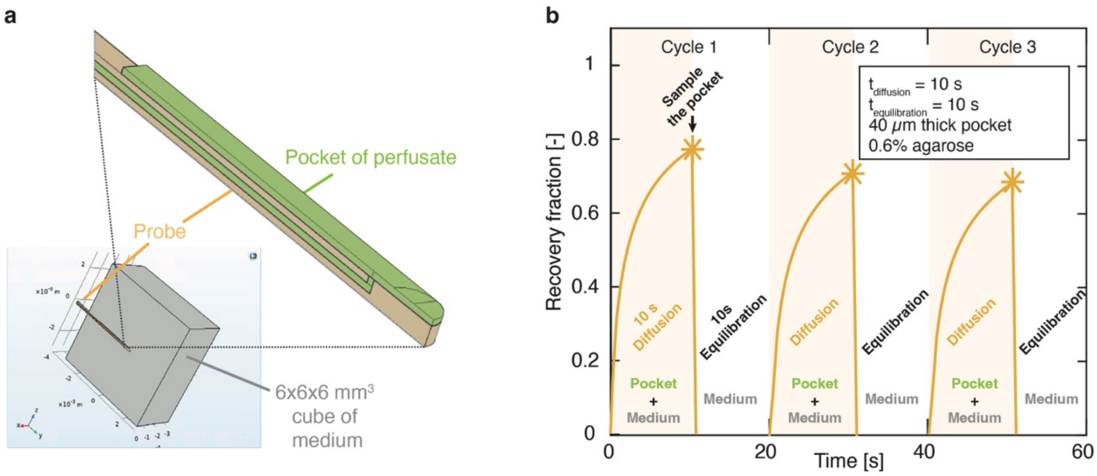


***Figure S2*** *The DoD model for simulations.* ***a*** *The model considers the probe at the center of a cube of 0.6% agarose, loaded with an initially uniform concentration of glucose (1 µM), and with a flat pocket of perfusate (0 µM) between the probe and the medium. Diffusion is the only mass transport phenomenon.* ***b*** *Average recovery fraction in the pocket of perfusate over three simulated DoD cycles. The 10 s long diffusion step considers diffusion in the medium and the pocket, where analytes diffuse to the pocket. At the time of sampling, at the end of the diffusion step, the average recovery fraction in the droplet is evaluated and marked as a star. The pocket is then removed from the model and the 10 s equilibration step considers diffusion in the medium only, to equilibrate the tissue concentration. At the beginning of the next cycle, a new pocket is considered (0 µM), but the concentration of the medium is given by the state of the medium at the end of the previous cycle.*

### FLOW ACTUATION DIAGRAM

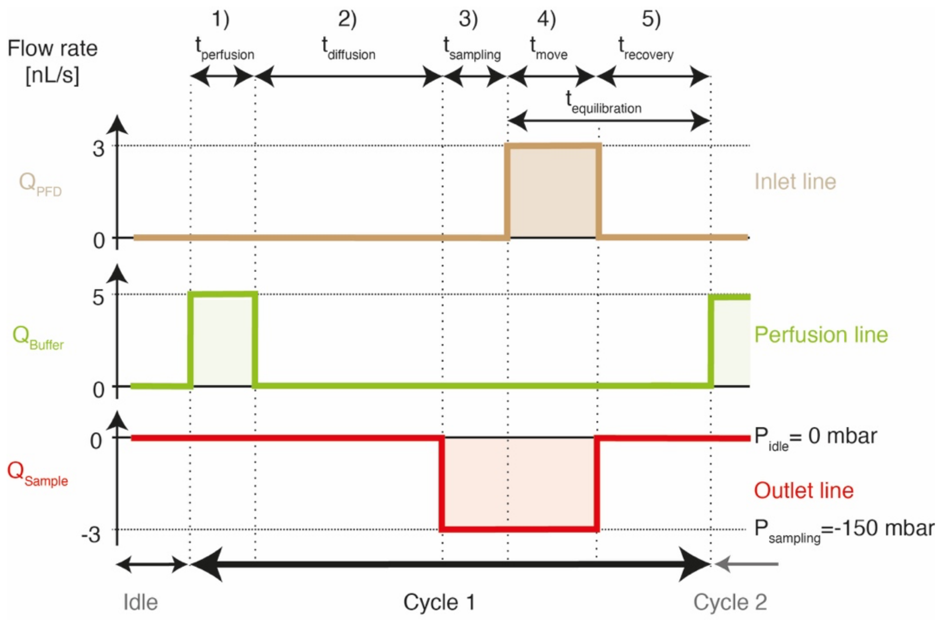


***Figure S3*** *Flow actuation diagram for the inlet, perfusion and outlet lines over a DoD cycle. All flows are at 0 nL/s in idle mode. The cycle 1 starts with the infusion of a controlled volume of perfusate in the tissue on the perfusion line only, creating a pocket of perfusate (1). The diffusion step occurs with all lines at 0 nL/s and molecules from the tissue diffuse to the pocket (2). The sample is aspirated from the tissue by pulling the outlet line to negative pressure during the sampling step (3). PFD is circulated inside the probe from the inlet line, to move the sample to the outlet line and the net flow to the tissue is null (4); all lines are put back to 0 nL/s for the recovery step until the next sampling cycle (5).*

### DOD SAMPLING IN VITRO – CONCENTRATION OF SUBSEQUENT DOD SAMPLES

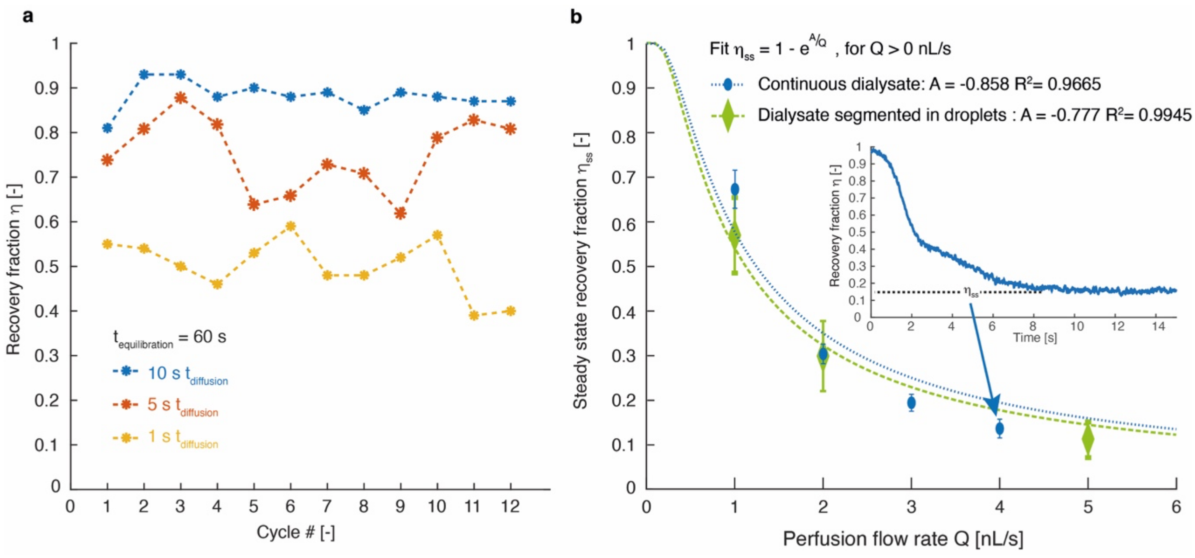


***Figure S4*** *Recovery fraction of NaCl sampled from 0.6% agarose by DoD and continuous sampling.* ***a*** *The recovery fraction is displayed for droplets subsequently sampled by DoD, with a fixed equilibration time and varied diffusion times of 1, 5 and 10 s. Each droplet is marked as a star. The dashed line is a visual guide showing the absence of transient regime.* ***b*** *The steady state recovery fraction of the sampled fluid is displayed, using the probe for continuous push-pull sampling, for different perfusion flow rates. This was accomplished by continuously infusing the gel with perfusate, which was directly sampled back at the outlet. The green series (diamonds) reports the case where dialysate was segmented into droplets, whereas the dialysate was continuous in the blue series (circles). The dashed lines were fitted to the experimental data, using a characteristic function for the recovery fraction in microdialysis[1]. The inset shows the instantaneous recovery fraction and the short transient regime when continuous push-pull sampling is started at 4 nL/s.*

### DOD SAMPLING IN VIVO – ELECTRICAL DROPLET DETECTION

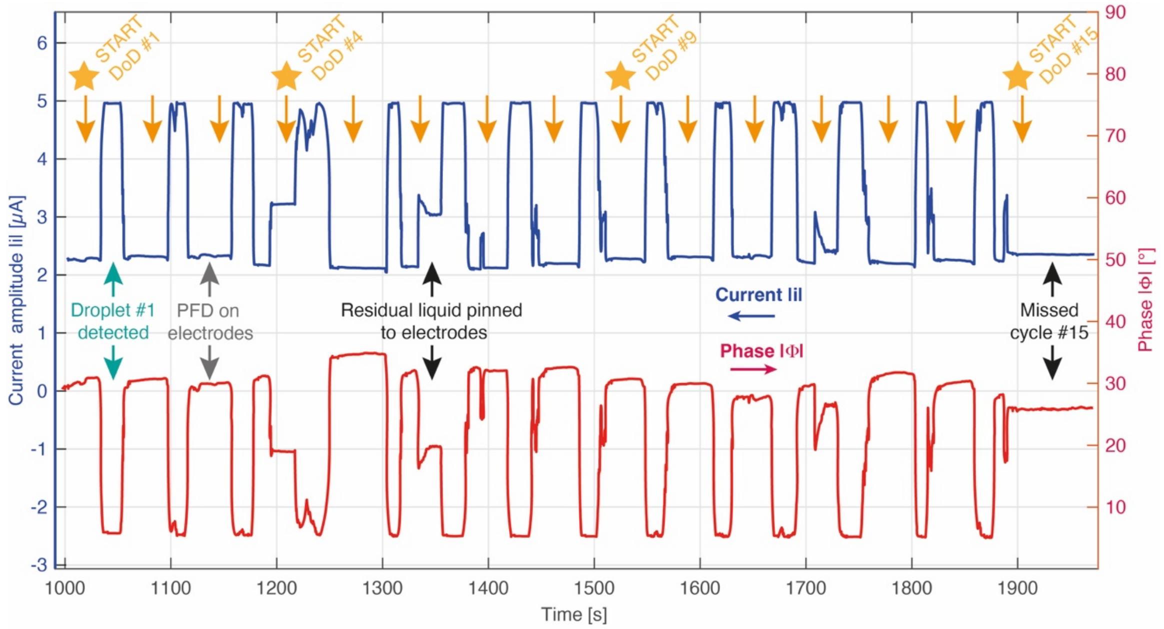


***Figure S5*** *Electrical detection of sampled droplets in vivo by measuring the current (blue, left axis) and the phase (red, right axis) across the electrodes in the outlet channel, using 100 mV excitation at 1 MHz. 15 DoD cycles were executed with 61 s interval. Cycles 1 to 14 successfully generated a droplet, whereas cycle 15 missed. Residual aqueous fluid pinned to the electrodes can be discriminated from droplets based on the current amplitude and on the phase at intermediate levels.*

### DOD SAMPLING IN VIVO – FLOW RATE

Figure S6 reports the pressure and the flow rate in the outlet line during two DoD cycles *in vivo*. The readout of the flow sensor enables to ensure that correct sampling parameters are used, not causing ultrafiltration, and matching the outlet flow rate to the flow rate of PFD during the droplet moving step. Figure S6a reports a fast sampling cycle. The perfusion step starts in (i), and no flow is expected in the outlet line. The apparent positive flow rate is an artifact from the pressure change at the end of the move step from the previous cycle, due to compliances in the system (air bubbles, etc.). This artifact disappears fast, and the outlet flow rate then correctly reads 0 nL/min until the end of the diffusion step (ii). Sampling starts by pulling the pressure to -300 mbar and another compliance artifact appears (iii) until the actual sampling flow rate is reached (iv). When the buffer pocket is fully sampled, the flow rate changes (v) and goes back to 0 nL/min due to the high hydraulic resistance of the tissue (vi). At the beginning of the droplet moving step (vii), PFD is flowed at the inlet at 300 nL/min and collected to the outlet, of which the flow rate goes back to -300 nL/min until the end of the moving step (viii). The pressure is set back to 0 mbar for the recovery step and another compliance effect (ix) appears prior to the start of the next DoD cycle. Figure S6b reports the outlet flow rate pattern during another DoD cycle, where the pressure and the duration of the sampling phase were adjusted to account for the change of hydraulic resistance of the system due to the accumulation of droplets in the capillary. With the adjusted sampling parameters, the transition from the sampling phase to the droplet moving phase occurs seamlessly. The states (v), (vi) and (vii) do not appear anymore. No ultrafiltration occurs and no PFD is dispensed into the tissue.

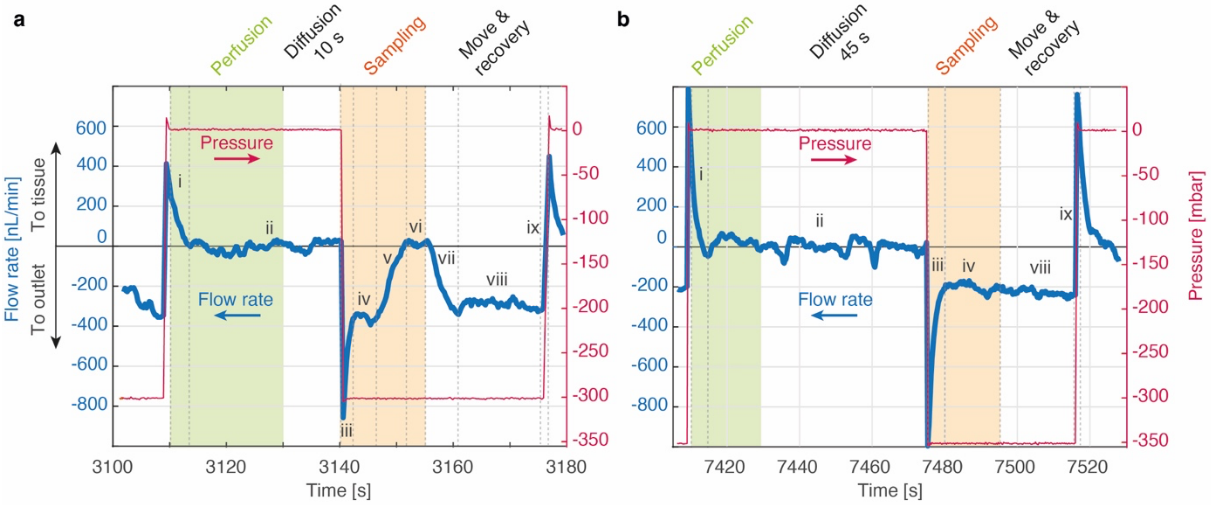


***Figure S6*** *Pressure actuation (right axis) and resulting flow rate (left axis) in the outlet line for two DoD cycles applied in vivo at different stages of a sampling experiment.* ***a*** *A cycle is applied, with 60 nL perfusion over 20 s, 10 s diffusion, 15 s sampling, 20 s droplet moving and 1 s recovery. The unoptimized sampling parameters lead to fast droplet sampling (iv) and potential ultrafiltration (vi).* ***b*** *A cycle is applied with 60 nL perfusion over 20 s, 45 s diffusion, 20 s sampling, 20 s droplet moving and 1 s recovery. The parameters are optimized for sampling and droplet moving without ultrafiltration or PFD dispense into the tissue.*

**DOD SAMPLING IN VIVO – PARAMETERS OF MOVIE OF 6 DOD CYCLES IN VIVO**

The Supporting Movie S1 (see MP4 file) shows the droplets collected at the outlet of the probe, over 6 DoD cycles with 60 nL perfusion over 20 s, 10 s diffusion, 15 s sampling, 20 s moving + 2 s recovery. Cycles 1 and 2 happen at real speed, cycles 3-6 happen at 10 times the real speed.

**DOD SAMPLING PARAMETERS**

*Table S1 Sampling parameters for experiments*

| *Figure* | *Perfusion volume* [nL] | *Diffusion time* [s] | *Sampling time* [s] | *Equilibration time* [s] | *Number of samples* [-] |
|---|---|---|---|---|---|
| *Fig. 3d* | 30 | 1 | 10 | 60 | 17 |
| | 30 | 3 | 10 | 60 | 22 |
| | 30 | 5 | 10 | 60 | 28 |
| | 30 | 10 | 10 | 60 | 35 |
| | 30 | 30 | 10 | 60 | 17 |
| | 30 | 60 | 10 | 60 | 13 |
| *Fig. 5a* | 30 | 10 | 20 | 30 | 10 |
| *Fig. 5b* | 30 | 1 | 20 | 30 | 45 |
| | 30 | 5 | 20 | 30 | 26 |
| | 30 | 10 | 20 | 30 | 17 |
| *Fig. 5b* | 60 | 1 | 20 | 30 | 13 |
| | 60 | 10 | 20 | 30 | 11 |
| | 60 | 30 | 20 | 30 | 11 |
| | 60 | 60 | 20 | 30 | 6 |

**REFERENCES FOR SUPPORTING INFORMATION**


1. Bungay, P. M., Morrison, P. F., Dedrick, R. L., Chefer, V. I. & Zapata, A. Chapter 2.2 Principles of quantitative microdialysis. in *Handbook of Behavioral Neuroscience* vol. 16 131–167 (2006).
